# SGK1.1 activation causes early seizure termination via effects on M-current

**DOI:** 10.1101/520775

**Authors:** Natalia Armas-Capote, Laura E. Maglio, Leonel Pérez-Atencio, Elva Martin-Batista, Antonio Reboreda, Juan A. Barios, Guadalberto Hernandez, Diego Alvarez de la Rosa, José Antonio Lamas, Luis C. Barrio, Teresa Giraldez

## Abstract

Early termination of *status epilepticus* affords protection against brain damage and associated pathologies. Regulation of Kv7.2/7.3 potassium channels, underlying the neuronal M-current, is key for seizure control. This conductance is maintained during initiation of action potentials, affecting neuronal excitability and thus inhibiting epileptic discharges. The M-current is upregulated by the neuronal isoform of the serum and glucocorticoid-regulated kinase SGK1 (SGK1.1). We tested whether SGK1.1 could act as an anticonvulsant factor using the kainic acid (KA) model of acute seizures in a transgenic mouse model with expression of a constitutively active form of the kinase. Our results demonstrate that SGK1.1 confers robust protection against seizures associated to lower mortality levels, independently of sex or genetic background. SGK1.1-dependent protection results in reduced number, shorter duration, and early termination of EEG seizures. At the cellular level, it is associated to increased M-current amplitude mediated by Nedd4-2 phosphorylation, leading to decreased excitability of hippocampal CA1 neurons without alteration of basal synaptic transmission. Altogether, our results reveal that SGK1.1-mediated M-current upregulation in the hippocampus is a key component of seizure resistance in the KA epileptic paradigm, suggesting that regulation of this anticonvulsant pathway may improve adverse outcomes to *status epilepticus*, constituting a potential target for antiepileptic drugs.

## Introduction

*Status epilepticus* (SE) is a life-threatening emergency characterized by a state of continuous behavioral or electroencephalographic (EEG) seizures (1, 2) and is considered one of the initial insults by which a normal brain becomes epileptic (3). Temporal Lobe Epilepsy (TLE) is one of the most common forms of epilepsy and the most refractory to treatment in humans. It originates in restricted limbic structures (temporal lobe: hippocampus, parahippocampal areas and amygdala). Long-term effects of this challenge depend on prompt seizure termination that will secure brain protection against permanent neuronal damage and associated pathologies (4).

In neurons, Kv7.2 and Kv7.3 (KNCQ2 and KCNQ3 genes) form heterotetrameric channels that constitute the molecular basis of the neuronal M-current, named after its well-known inhibition via muscarinic receptor activation, which underlies some cholinergic excitatory pathways (5, 6). Kv7.2/7.3 activation counteracts brain excitability since they are active at subthreshold potentials near to −60 mV, and do not inactivate under persistent depolarization (7). This leads to a membrane potential stabilization under subthreshold excitatory inputs, preventing repetitive action potential firing (6). Thus, Kv7.2/7.3 channels constitute a potent inhibitory mechanism that is especially relevant in hyperexcitability disorders such as epilepsy. We previously described a novel physiological mechanism of M-current regulation via activation of SGK1.1, the neuronal isoform of the serum and glucocorticoid-regulated kinase 1 (SGK1) (8). This isoform arises from alternative splicing of the *Sgk1* gene (9). The ubiquitous and neuronal isoforms share the catalytic and activation domains but differ in their N-terminal domain. SGK1.1 shows enhanced stability and is tethered to the plasma membrane by binding to phosphatidylinositol 4,5-bisphosphate (PIP_2_) (9, 10), a primary controller of M-current activity (11). The SGK1.1 regulatory pathway provides a potential confluent point between muscarinic-dependent inhibition of Kv7.2/7.3 channels and their SGK1.1-mediated upregulation (8).

Using a transgenic mouse model (B6.Tg.sgk1) generated by the introduction of a modified bacterial artificial chromosome harboring the full-length *Sgk1* gene with an activating point mutation leading to increased SGK1 activity, our previous work hinted that SGK1.1 confers resistance to kainic acid (KA)-induced seizures by upregulating the M-current in the brain (8). We now demonstrate that SGK1.1 activity confers resistance to KA-seizures, potently decreasing seizure severity. Electroencephalographic analysis revealed that SGK1.1 associates with early termination of seizure events, which in addition occur at significantly lower number and shorter durations. These effects depend on upregulation of the M-current via Nedd4-2 phosphorylation, leading to reduced hippocampal neuronal excitability without affecting basal synaptic transmission. Taken together, these results support a relevant role of SGK1.1 in regulating neuronal excitability as an anticonvulsant factor that may improve deleterious outcomes of SE.

## Results

### B6.Tg.sgk1 mice show reduced seizure severity and no lethality during acute KA-induced SE

Seizure behaviour after intraperitoneal injection of KA in B6.WT and B6.Tg.sgk1 mice was evaluated using a modified Racine scale (see Methods and (8)). Progression to stage 4, a more severe and convulsive stage, was significantly reduced in B6.Tg.sgk1 mice when compared to B6.WT (**Figure 1A**). Around 60% B6.WT showed tonic-clonic seizures (stage 6) compared to 17% B6.Tg.sgk1 mice. In consistence with previous reports (12), 20% B6.WT mice reached stage 7 (death). Strikingly, mice did not display lethality events **(Figure 1A)**.

**Figure 1.**
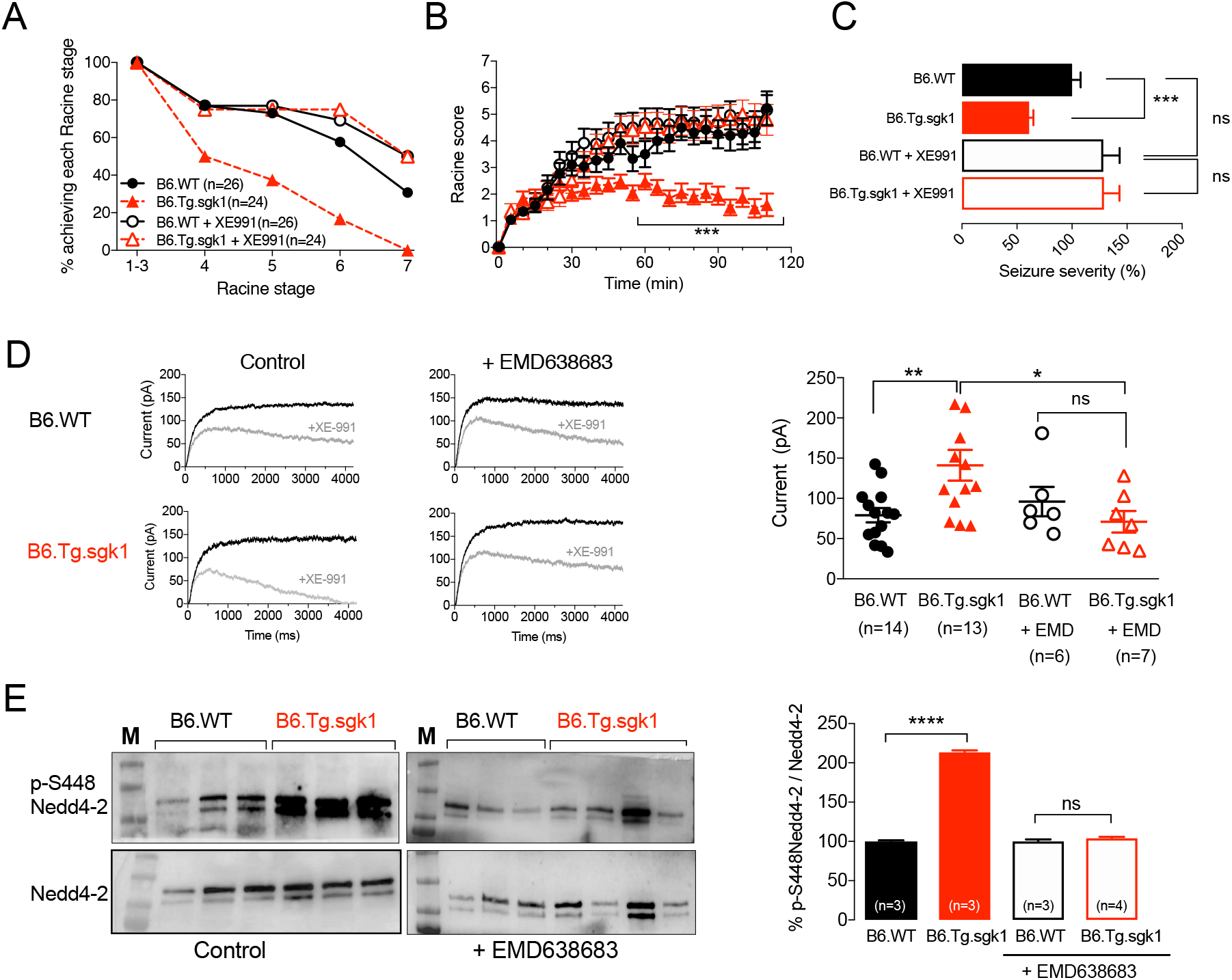
B6.Tg.sgk1 mice show reduced seizure severity after systemic KA administration via M-current activation. (**A)** Cumulative plot representing the percentage of mice reaching the indicated Racine stages. Number of experiments for each condition is indicated in the legend. (**B)** Raw Racine stage score (mean ± SEM) in 5 min intervals over the monitored period of time. Symbols and number of experiments are the same as in legend of panel A. *** p < 0.002, *two-tailed Student t-test*. (**C)** KA-induced seizure severity values for both mice genotypes, with and without previous treatment with XE991. Values are mean ± SEM. *** p < 0.0005, *ns* not significant, *two-way ANOVA*. **(D)** *Left*, Representative current traces in response to a depolarizing pulse from −70 mV to −30 mV before (black) and after addition of 3 μM XE991 (gray), in the absence and presence of 50 μM EMD638683. *Right*, M-current values for each group of neurons without and with pre-incubation with EMD638683. * p < 0.05; ** p < 0.01, n.s. non-significant, *two-way ANOVA*. **(E)** *Left*, Representative immunoblots for detection of total Nedd4-2 (*lower panels*) and its phosphorylated form (P-S448; *upper panels*) in hippocampus lysates from B6.WT and B6.Tg.sgk1, without and with pre-treatment with EMD638683. *Right*, Quantification of P-S448-Nedd4-2 (relative to total Nedd4-2). **** p < 0.0001, *ns* not significant, *Unpaired Student-test*.

Evaluation of the time course of KA-induced seizure behavior by scoring Racine stage averages in 5 min intervals throughout the experiment (13) revealed an acute progression towards higher Racine scale stages during the first 30 min after KA administration in both genotypes **(Figure 1B)**. However, after this initial period B6.WT mice progressively reached more severe seizure behavior stages, while transgenic mice stabilized and subsequently showed a reversion pattern **(Figure 1B)**. We integrated the individual scores per mouse for the total duration of the experiments to better account for seizure severity in mice that died during the experiment, as described previously (13). As shown in **Figure 1C**, seizure severity was significantly reduced in B6.Tg.sgk1 mice. Importantly, the protective effect of SGK1.1 activation against seizures was independent of sex and genetic background, which have been shown to affect seizure susceptibility in various animal paradigms (12, 14-19) (**Supplemental Figure 1**).

Altogether, our results demonstrate that SGK1.1 activation in transgenic mice confers less susceptibility to KA-induced behavioral seizures and reduced mortality. Additionally, they suggest that the transgene affects the time course of SE evolution.

### Reduced seizure severity in B6.Tg.sgk1 mice depends on M-current activation

Isolated superior cervical ganglion (SCG) sympathetic neurons from B6.Tg.sgk1 show enhanced M-current activity paralleled with decreased excitability and more negative resting potentials (8). We addressed the role of M-current upregulation in resistance to KA-induced seizures and reduced mortality after *Sgk1* activation by treating mice with a specific M-current inhibitor, XE991. As shown in **Figure 1A-B**, the protective effect of *Sgk1* activation was abolished in transgenic mice pre-treated with XE991. Accordingly, no differences in seizure severity were observed in WT *vs.* transgenic mice after treatment with the inhibitor (**Figure 1C**). These effects on seizure behavior and M-current regulation were not associated to alterations in the gene expression of KA receptors (GluK1-5; GRIK1-GRIK5 genes) or Kv7.2/3 channels caused by constitutively active *Sgk1* in the B6.Tg.sgk1 transgenic mice, as demonstrated by quantitative PCR (qPCR) experiments showing significant alterations in KCNQ2, KCNQ3 or GRIK1-5 mRNA expression in the hippocampus (**Supplemental Figure 2**).

Altogether, these data strongly suggest that upregulation of M-current function constitutes the mechanism underlying protection against convulsions driven by *Sgk1* activation.

### CA1 pyramidal neurons from B6.Tg.sgk1 showed increased M-current amplitude

The effect of SGK1.1 in relevant epileptogenic areas of the brain, including hippocampus (20), remain unknown. We performed whole-cell patch-clamp recordings in CA1 pyramidal neurons (CA1-PN) from B6.Tg.sgk1 and B6.WT mice hippocampal slices. B6.Tg.sgk1 CA1-PN showed significantly higher levels of M-current, measured as the XE991-sensitive current (**Figure 1D**). Importantly, this difference was abolished after addition of the SGK1 inhibitor EMD638683 **(Figure 1D**, *right panel*). These results support the idea that SGK1.1-mediated upregulation of the M-current in CA1-PN underlies the seizure-resistant phenotype observed in Tg.sgk1 mice.

In heterologous expression systems, activation of *Sgk1* modulates M-current amplitude by increasing Kv7.2/7.3 channels cell surface expression through a mechanism involving the ubiquitin ligase Nedd4-2 (8, 21, 22). We observed a significant increase in phosphorylation levels of Nedd4-2 in HEK293 cells transiently co-transfected with SGK1.1 and Nedd4-2 (**Supplemental Figure 3A**). Additionally, phosphorylation of endogenously-expressed GSK3β, another known target of SGK1 (23), was increased when SGK1.1 was transiently transfected in HeLa cells (**Supplemental Figure 3B**), suggesting that SGK1 isoforms share phosphorylation targets. Importantly, in B6.Tg.sgk1 hippocampal tissue we observed higher phosphorylation of Nedd4-2, which was prevented after incubation with the SGK1 inhibitor EMD638683 (**Figure 1E-F**). Our data demonstrate that Nedd4-2 is a substrate of SGK1.1 in hippocampal neurons, constituting a regulatory pathway underlying M-current modulation by this kinase in hippocampus.

### Epileptic EEG activity during KA-induced SE is reduced in B6.Tg.sgk1 mice

Video recordings coupled to EEG allowed us to investigate the neurophysiological basis underlying the different time course of behavioral Racine stages between transgenic and WT mice. **Figure 2** shows representative recordings of hippocampal and cortex EEG activity in B6.Tg.sgk1 and B6.WT adult male mice, 1 h before and up to 6 h after KA injection. Evolution of epileptic activity was quantitatively evaluated using time–frequency analysis and the normalized amplitude of hippocampal and cortex signals. Following KA injection, B6.WT mice responded initially (0-30 min) with seizure activity of increasing amplitude and power of multiple frequencies reaching the first generalized seizures (GS**; Figure A-1 and Ci-1)** followed by postictal depression. SE worsened over the following 1-2 h with increasing number of GS and interictal spiking activity **(Figure A-2 and C-i2)**, after which GS rate decreased progressively. An intensive spiking activity still remained within the 6^th^ hour of recording **(Figure A-3 and C-*i*3)**. In contrast, B6.Tg.sgk1 mice showed significantly lower SE severity; although SE progressed initially in a similar manner (0-30 min), a more rapid and profound reduction in the number of GS and spiking activity occurred over time. Seizures subsided and the EEG returned to normal activity within 6 h of recording (**Figure 2B and 2Cii)**.

**Figure 2.**
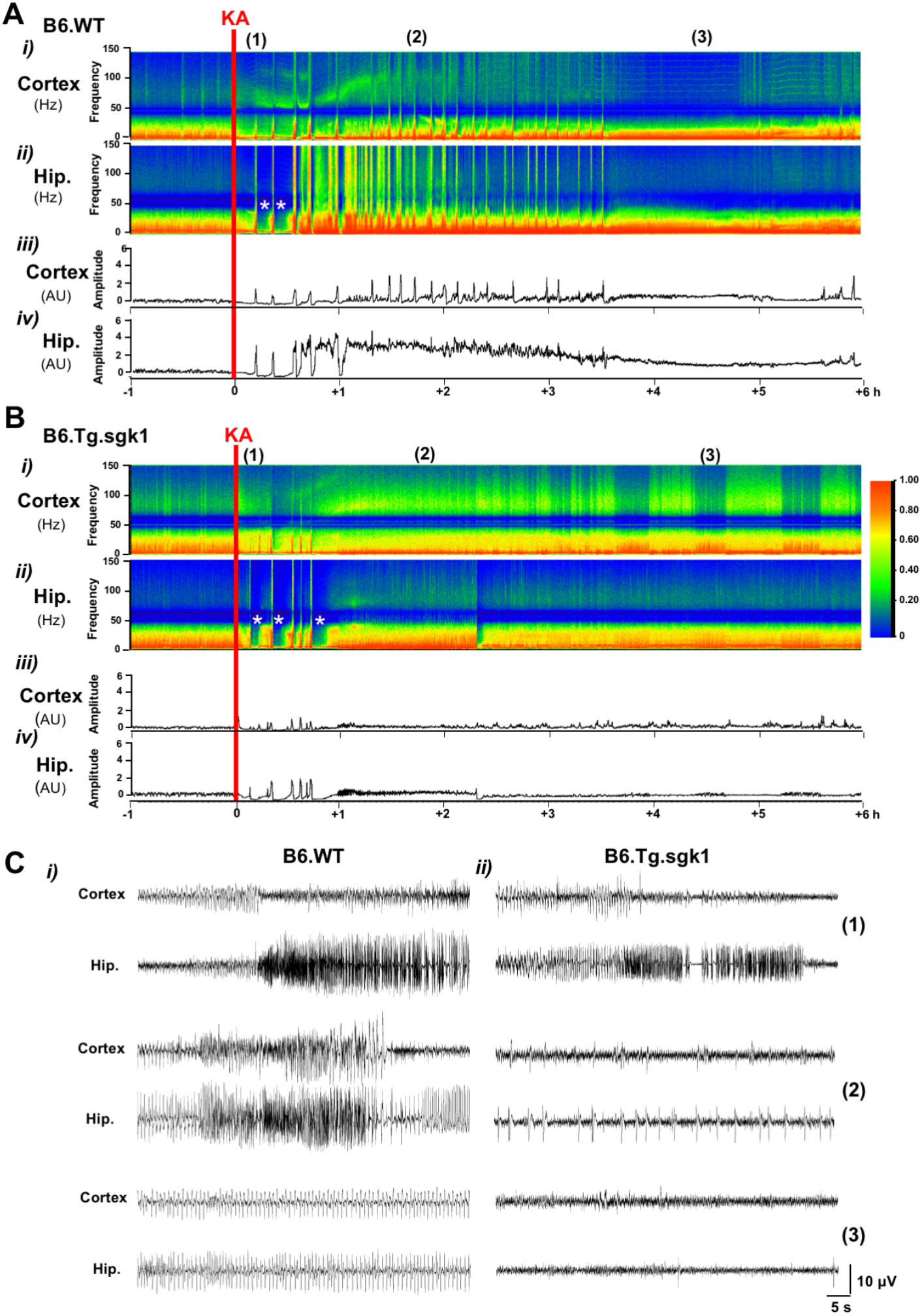
EEG recordings show reduced severity of KA-induced seizures in B6.Tg.sgk1 mice. Time-frequency plots of cortex and hippocampus signals (*i* and *ii*) and the corresponding normalized values of amplitude (*iii* and *iv*) 1 h prior to and 6 h after KA injection (red line) in **(A)** B6.WT and **(B)** B6.Tg.sgk1 mice. Asterisks (*) denote postictal depression periods. **(C)** Insets of raw EEG traces from B6.WT mice (*i*) and B6.Tg.sgk1 mice (*ii*) at time points (1) to (3) indicated on top of panels A and B.

### Increased SGK1.1 activity reduces the number and duration of generalized seizures in hippocampus and cortex

GS number was significantly reduced in hippocampus and cortex of B6.Tg.sgk1 mice **(Figure 3A)**. This was appreciable after the first hour post-KA, when transgenic mice reverted towards a lower number of GS that practically disappeared within 3 hours. In contrast, B6.WT mice maintained significantly elevated GS numbers during at least 4 hours (**Figure 3A**).

**Figure 3.**
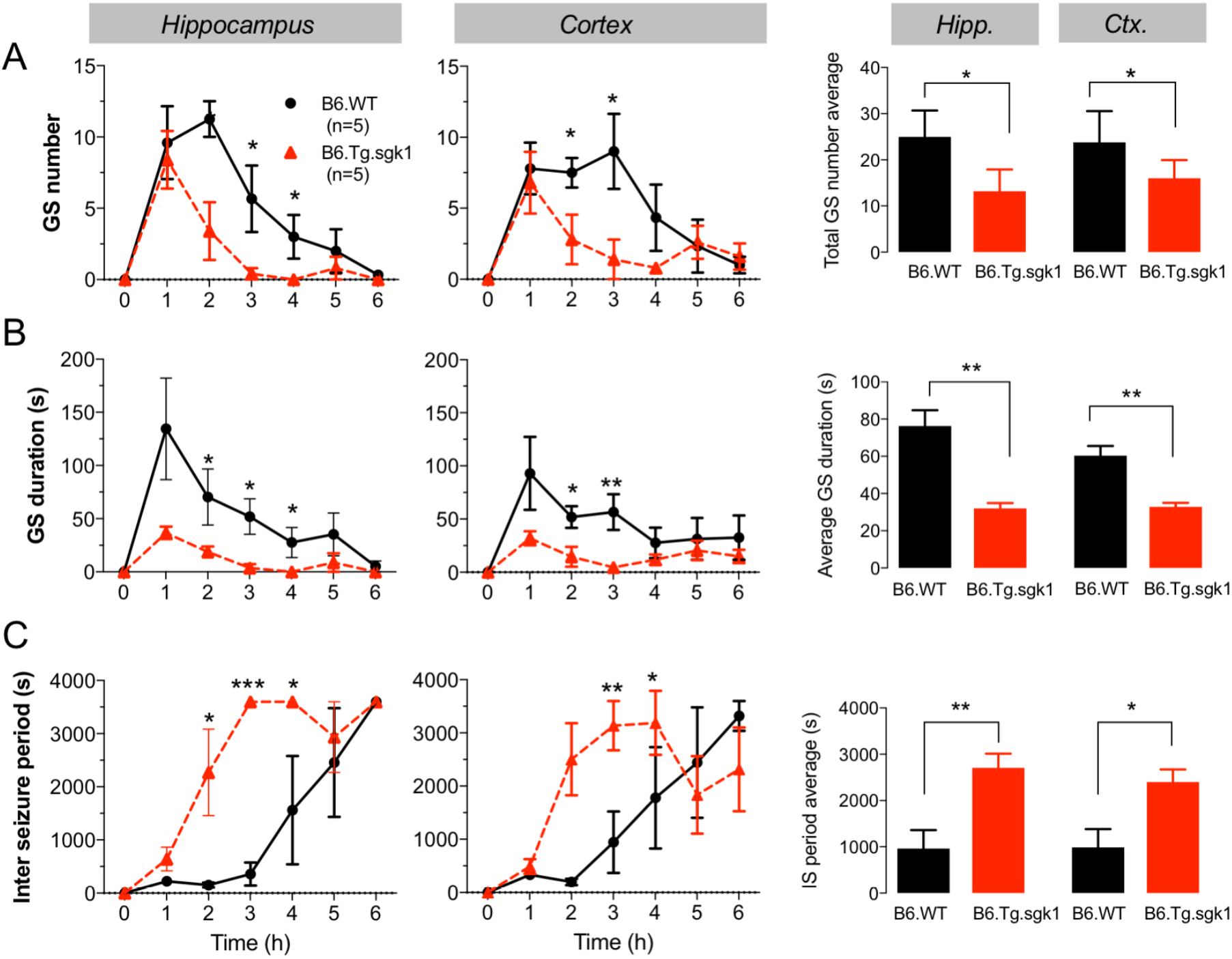
Activation of SGK1.1 reduces generalized seizure (GS) number and duration in hippocampus and cortex. (**A)** Time course of average number of GS after KA administration in hippocampus *(left)* and cortex *(middle)* and average GS number during the duration of the experiment (6 h) *(right)*. Data are presented as mean ± SEM corresponding to 1 h intervals; * p < 0.05, *two-tailed Student t-test*. **(B)** Time course of GS duration after KA administration in hippocampus *(left)* and cortex *(middle)*, and average GS duration corresponding to the duration of the experiment *(right)*. Data are presented as mean ± SEM corresponding to 1 h intervals; * p < 0.05, ** p < 0.01, *two-tailed Student t-test*. **(C)** Time course of inter-seizure periods after KA administration in hippocampus *(left)* and cortex *(middle)* and average inter-seizure period during the experiment (6 h) *(right)*. Data are presented as mean ± SEM corresponding to 1 h intervals; * p < 0.05, ** p < 0.01, *** p < 0.001, *two-tailed Student t-test*.

In addition to the number of GS, seizure duration is a key factor affecting the recovery from acute SE that has been highlighted as a determinant for brain damage and mortality (24). EEG experiments revealed significantly lower seizure duration in B6.Tg.sgk1 mice **(Figure 3B)**. We also observed a significant increase in the inter-seizure period for B6.Tg.sgk1 mice within 2-4 hours after KA administration (**Figure 3C**), indicating a faster time course for seizure termination. This result is consistent with the observed lower lethality levels (**Figure 1**) and diminished brain damage in transgenic mice (unpublished observations). In summary, our results demonstrate that the SGK1.1 anticonvulsant effect involves reduction of both the number and the duration of epileptic events, as well as an earlier termination of seizure activity.

### Basal synaptic transmission is not modified by increased SGK1.1 activity

EEG results prompted us to study the effect of *Sgk1* activation on neuronal excitability in hippocampus. First, we characterized basal synaptic transmission by recording miniature excitatory or inhibitory postsynaptic currents (mEPSC and mIPSC) from hippocampal CA1-PN. Constitutive activation of SGK1.1 in B6.Tg.sgk1 mice had no effect on the frequency of mEPSC (**Figure 4A-B**) and mIPSC (**Figure 4E-F**), suggesting that *Sgk1* activation does not modulate the probability of glutamate or GABA release at the synapse. Additionally, no differences were observed on the amplitude of mEPSC (**Figure 4C-D**) nor mIPSC (**Figure 4G-H**) recorded from B6.Tg.sgk1 *vs.* B6.WT, suggesting that *Sgk1* activation has no significant effect on the functional levels of AMPA/NMDA (mEPSC) nor GABA_A_ (mIPSC) postsynaptic receptors.

**Figure 4.**
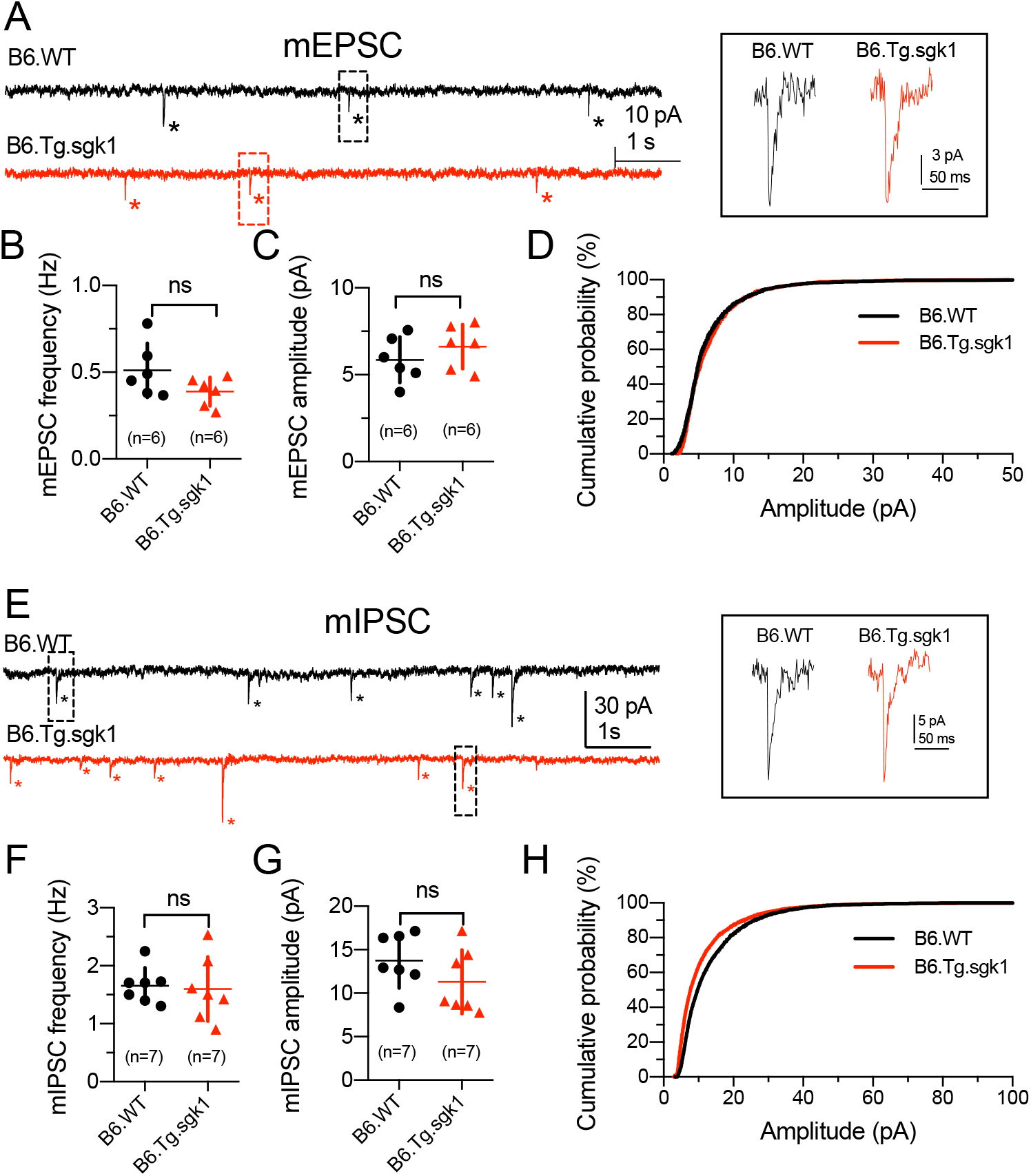
Basal synaptic transmission is not altered in B6.Tg.sgk1 mice. **(A)** Representative current traces recorded at −70 mV in CA1-PN from B6.WT *(black)* and B6.Tg.sgk1 *(red)* mice in the presence of 1 µM TTX, 50 µM PiTX, and 5 μM CPG-55845 to isolate mEPSCs. Asterisks denote mEPSC events. Insert shows representative mEPSCs in an expanded scale. **(B)** Frequency, amplitude, and **(D)** cumulative probability distribution of amplitudes of mEPSC in B6.WT *(black)* and B6.Tg.sgk1 mice *(red)*. **(E)** Representative current traces recorded at −70 mV in CA1- PN from B6.WT *(black)* and B6.Tg.sgk1 *(red)* mice in the presence of 1 µM TTX, 50 µM AP5, and 20 μM CNQX to isolate mIPSCs. Asterisks denote mIPSC events. Insert shows representative mIPSCs in an expanded scale. **(F)** Frequency, **(G)** amplitude, and **(H)** cumulative probability distribution of amplitudes of mIPSC in B6.WT *(black)* and B6.Tg.sgk1 mice *(red). ns*, not significant*, unpaired two-tailed t-test*.

### B6.Tg.sgk1 CA1 pyramidal neurons show reduced excitability

In SCG neurons, the M-current regulates resting membrane potential and some relevant features of neuronal function (7, 25). We assessed the influence of *Sgk1* activation on hippocampal CA1-PN intrinsic membrane properties and action potential characteristics by performing whole-cell current-clamp recordings in tissue slices. Interestingly, we found no significant differences between genotypes (**Supplemental Figure 4**), indicating that passive and active properties of CA1-PN are unaffected by the expression of the transgene.

The M-current is considered a key component in regulatory mechanisms of excitability by controlling spiking frequency, among other mechanisms (6, 26, 27). Therefore, we determined whether CA1-PN from B6.Tg.sgk1 mice show altered excitability, defined as the overall tendency to generate evoked action potentials in response to increasing current injection steps. We observed a significant reduction of firing frequency in B6.Tg.sgk1 CA1-PN **(Figure 5A-B**), indicating reduced excitability in transgenic neurons. We next evaluated KA-induced alterations in membrane properties and excitability in hippocampal CA1-PN. Application of KA induced a significantly smaller increase in neuronal firing frequency in B6.Tg.sgk1 than in B6.WT mice (**Figure 5B**). Additionally, the change in membrane potential towards more depolarized values associated to KA treatment (28) was significantly smaller in B6.Tg.sgk1 compared to B6.WT neurons (**Figure 5C)**, suggesting that constitutively active SGK1.1 induces a resistance to depolarization induced by KA hippocampal CA1 neurons.

**Figure 5.**
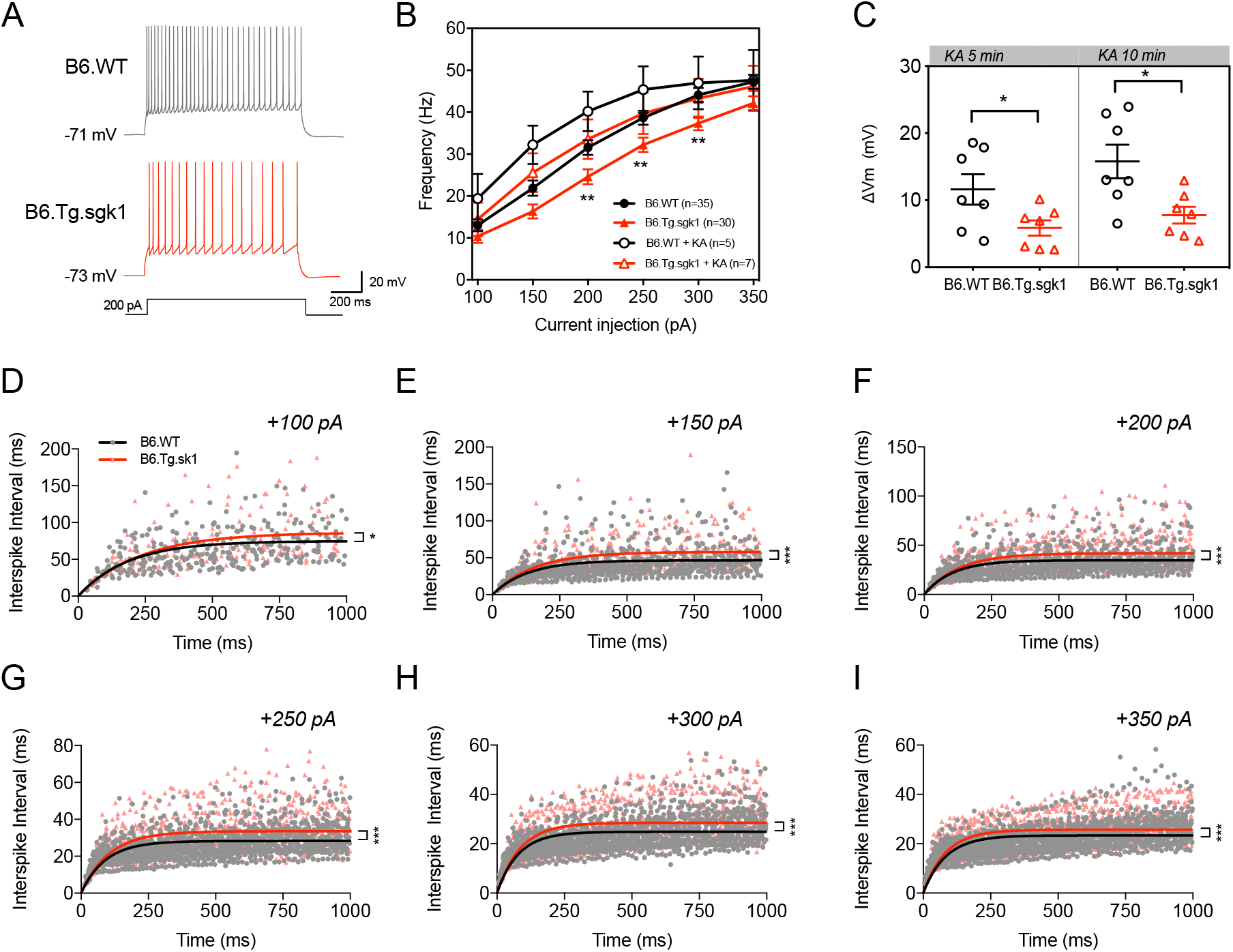
Intrinsic excitability is reduced in CA1-PN from B6.Tg.sgk1 mice. **(A)** *Left*, representative traces recorded in CA1-PN after a 200 pA current injection in B6.WT *(black)* and B6.Tg.sgk1 *(red)* mice. **(B)** Frequency-current relationships for CA1-PN from B6.WT *(black)* and B6.Tg.sgk1 *(red)* mice in the absence *(full symbols)* and the presence *(open symbols)* of KA; ** p < 0.01, *Multiple t-test*. **(C)** Resting membrane potential increments 5 min *(left)* and 10 min *(right)* after KA application in CA1-PN from B6.WT *(black)* and B6.Tg.sgk1 *(red)* mice; * p < 0.05, *unpaired Student t-test*. **(D-I)** CA1-PN interspike intervals *vs* time after current injections indicated on top of graphs, from B6.WT *(black)* and B6.Tg.sgk1 *(red)* mice. Solid lines correspond to exponential function fit to the data; * p < 0.05, *** p < 0.001, *F-test*.

Spike frequency adaptation (SFA) is an intrinsic neuronal property that depends on the specific battery of ion channels expressed at the membrane of the different neuronal types. SFA is characterized by a progressive increment in the interspike intervals (ISIs) (29). It has been demonstrated that M-current activation during neuronal firing regulates ISIs, leading to increased SFA in many neuronal types, including CA1-PN (30-32). In B6.Tg.sgk1 CA1-PN, values of CA1- PN ISIs over time reached a plateau that was significantly higher than that observed in B6.WT neurons (*e.g*., 42.0±0.8 ms in B6.Tg.sgk1 *vs.* 34.8±0.5 in B6.WT at 200 pA current-injection; p<0.001**; Figure 5F)**, indicating that activation of SGK1.1 may be controlling SFA by regulating ISIs.

## Discussion

In this study, we have shown that transgenic expression of the *Sgk1* gene including a mutation for constitutive activation confers remarkable protection against seizures in the KA model. In neurons, *Sgk1* function mainly results in expression of the SGK1.1 isoform (9), which regulates membrane abundance of Kv7.2/7.3 tetramers underlying the M-current (8). The protection against KA-induced seizures was lost when animals were treated with the specific M-current inhibitor XE991, which also abolished the decrease in membrane excitability in transgenic CA1-PN. Altogether, these results demonstrate that the upregulation of M-current is the main mechanism underlying the effect of SGK1.1. The protection effect of *Sgk1* is robust, with little variation due to factors such as sex or genetic background, which commonly act as modifiers in the phenotypic effects of genetic manipulation. EEG recordings demonstrated that seizure protection in transgenic mice arises mainly from reduced number of GS, which additionally showed shorter duration, together with early SE termination.

Behavior analysis of KA-induced seizures showed clear differences in the temporal development of seizure patterns, indicating a possible effect on seizure progression. EEG recordings from KA-injected animals clearly showed that the onset of seizures is not altered. Strikingly, transgenic mice showed a significantly reduced number and shorter duration of seizure events in hippocampus and cortex, together with earlier termination of seizures. There are several mechanisms that could explain these observations. First, our results raise the question of whether transgenic mice present reduced KA bioavailability. Our finding that B6.WT and B6.Tg.sgk1 mice show comparable seizure latency values rules out the possibility that KA reaches the brain at lower concentration. Second, differential response to KA between genotypes could be due to variability in the amount or activity of GluK receptors. However, we observed no changes in mRNA expression of GRIK1-5 mRNA in hippocampus. Third, differences in the response to KA could be attributable to basal alterations in synaptic transmission (e.g., an increase or decrease in glutamatergic or GABAergic transmission) due to *Sgk1* activation. However, we demonstrate that both basal excitatory and inhibitory synaptic transmission are not different between both genotypes. Finally, a reduced effect of KA would be expected if transgenic animals showed intrinsically lower neuronal excitability. In light of our previous results obtained with M-current measurements in isolated SCG neurons from B6.Tg.sgk1 (8), it was reasonable to hypothesize that reduced excitability due to increased M-current magnitude plays a major role in resistance to KA-induced seizures. In fact, our results are reminiscent of the effects of retigabine, a Kv7.2/Kv7.3 channel opener that shortens number and duration of EEG spike bursts on KA-treated mice (33). This conclusion is reinforced by the effects of XE991 shown in this study, which revert the effect of KA injection in B6.Tg.sgk1 mice towards the B6.WT phenotype. Furthermore, our results demonstrate a reduction in the intrinsic excitability of CA1-PN from B6.Tg.sgk1, as reflected by a decrease in the firing frequency and the increase in SFA compared to B6.WT neurons under the same stimulation conditions.

In hippocampal CA1-PN, *Sgk1* activation was not associated to changes in intrinsic membrane properties or action potential characteristics. These results can be explained based on the fairly negative resting membrane potentials observed in hippocampal neurons, which are unlikely to be under the control of the M-current (that is activated between −60 mV and −20 mV, approximately). In these neurons, the role of the M-current will be made evident when the cell is depolarized (*e.g.*, after KA administration) or actively firing (therefore the observed effect on SFA (29, 30)).

One of the most remarkable results obtained in this study is the role of SKG1.1 on controlling the number and duration of epileptic events, playing an important role in limiting the progression of SE in the KA model. This is especially relevant not only to antagonize convulsive activity but also because early seizure termination has been connected to protection against permanent neuronal damage and associated pathologies (24, 34, 35). Moreover, it could explain the striking differences found in lethality rates between KA-treated transgenic and WT animals, particularly in adult male B6.Tg.sgk1, where there was no death. This is in accordance with high mortality rates associated to SE when seizure duration is longer than 1 h (24).

Drugs effective against KA-induced seizures have been proposed to be relevant for temporal lobe epilepsy (TLE) (20, 36). The potent anticonvulsant role of SGK1.1 against KA-induced seizures suggests that activation of SGK1.1 could be useful in seizure control during SE, which could in turn limit TLE progression. Further studies in the context of a chronic model of epilepsy such as pilocarpine treatment (37) will be necessary to demonstrate long-term effects on limiting TLE development.

## Material & Methods

### Animals

All experimental procedures were approved by the University of La Laguna, the University of Vigo, and the Hospital Ramón y Cajal Ethics Committees on Animal Welfare and conform to Spanish and European guidelines for the protection of experimental animals (RD1201/2005; 2010/63/EU). The B6.Tg.sgk1 mouse line was generated as previously described (8). FVB.Tg.sgk1 mice were generated by transferring the transgene to the inbred strain FVB/N, backcrossing for 9 generations. Wild type mice (B6.WT or FVB.WT) in the same genetic background (C57BL6/6J or FVB/N) were used as controls.

### Kainic acid administration and evaluation of seizure behavior

SE was induced by intraperitoneal injection of 20 mg/kg KA, as described previously (8). Where indicated, intraperitoneal injection of 10 mg/kg XE991 was performed 1 h prior to KA injection (38). Seizure behavior was scored using a modified Racine scale (39): immobility (*stage 1)*; forelimb and/or tail extension (*stage 2*); repetitive movements, head bobbing (*stage 3*); rearing and falling (*stage 4*); continuous rearing and falling, jumping and/or wild running (*stage 5*); generalized tonic-clonic seizures (*stage 6*); death (*stage 7*). Seizure severity was determined as previously described (13):

Seizure Severity = Σ (all scores of a given mouse)/time of experiment

The severity of all genotypes or conditions tested was normalized to WT mice (13).

### Brain slice preparation

Anesthetized 4-6 week old male mice (ketamine:xylazine cocktail, 60:5 mg/kg) were intracardially perfused with ice-cold choline-based artificial cerebrospinal fluid (ACSF-Ch), containing (in mM): 110 choline chloride, 1.25 NaH_2_PO_4_, 25 NaHCO_3_, 7 MgCl_2_, 0.5 CaCl_2_, 2.5 KCL, 7 glucose, 3 pyruvic acid, and 1.3 ascorbic acid. Coronal hippocampal slices (350 μM thick), obtained using a VT1000S vibratome (Leica), were equilibrated in ACSF (in mM: 125 NaCl, 1.25 NaH_2_PO_4_, 25 NaHCO_3_, 2 MgCl2, 1.6 CaCl_2_, 2.5 KCl, and 10 glucose; pH = 7.35) constantly bubbled with carbogen (95% O_2_, 5% CO_2_), for ≥ 1 h before recordings.

### Whole-cell patch-clamp recordings

Hippocampal CA1-PNs were recorded using the whole-cell current-clamp and voltage-clamp configurations of the patch-clamp technique using an Axopatch amplifier and pCLAMP software (Molecular Devices). Slices were visualized with an upright microscope (Zeiss Axoskop 2FS or Olympus BX51WI) and a near-infrared charged-coupled device camera. Recordings were performed at 34±1.5°C in a submerged recording chamber continuously perfused with ACSF. 0.5μM tetrodotoxin (TTX; Sigma) was added in voltage-clamp experiments. Patch pipettes (5-8 MΩ) were pulled (P-87 puller, Sutter Instruments) from borosilicate glass capillaries (Science Products) and filled with an intracellular solution containing (in mM): 120 K-gluconate, 10 HEPES, 0.2 EGTA, 20 KCl, 2MgCl_2_, 7 di-Tris-phosphocreatine, 4 Na_2_ATP, 0.3 Tris-GTP, and 0.1% biocytin (pH = 7.3). Sampling rate was 20 kHz with low-pass filtering at 5 kHz.

### Miniature postsynaptic current recordings

Miniature inhibitory postsynaptic currents (mIPSC) were recorded at −70 mV in ACSF in the presence of 1 µM TTX, 20 µM 7-nitro-2,3-dioxo-1,4-dihydroquinoxaline-6-carbonitrile (CNQX; Tocris), and 50 µM D-2-amino-5-phosphonovaleric acid (D-AP-5; Tocris) to isolate synaptic inhibitory transmission. Recording pipettes were filled with an internal solution containing (in mM): 113 K+-gluconate, 25 KCl, 10 HEPES, 0.2 EGTA, 4 Na_2_ATP, and 0.3 Na_2_GTP (pH 7.3, 320-340 mOsm). Miniature excitatory postsynaptic currents (mEPSCs) were recorded at −70 mV in ACSF in the presence of 1 μM TTX, 20 μM picrotoxin (PiTX), and 5 µM CGP-55845 to isolate synaptic excitatory transmission. Recording pipettes were filled with an internal solution containing (in mM): 130 K+-gluconate, 8 KCl, 10 HEPES, 0.2 EGTA, 4 Na_2_ATP, and 0.3 Na_2_GTP (pH 7.3, 310–330 mOsm). Pipette resistance was 5-7 MΩ. Synaptic currents were filtered at 1 kHz (low pass), digitized (Digidata 1550B, Molecular Devices), and acquired by the pCLAMP software (Molecular Devices) with a PC computer. MiniAnalysis software (SynaptoSoft Inc) and pCLAMP software were used for the analysis of the frequency and amplitude of mIPSCs and mEPSCs.

### Cell culture and transfection

HEK293 and HeLa cells were obtained from the American Type Culture Collection (ATCC) and cultured according to the supplier’s recommendations. A plasmid encoding the full-length Nedd4-2 was transiently transfected into HEK293 cells following previously described procedures (8).

### Western blot

The hippocampal formation was isolated from brain slices (350 μm thick) with or without SGK1 inhibitor EMD638683 (50 μM) and transferred to lysis buffer (Tris-Base 0.5 M, pH 7.4; SDS 10%; phosphatase and proteinase inhibitors). TENT buffer (Tris-EDTA-NaCl-Triton, plus phosphatase and proteinase inhibitors) was used for cultured cells. Total and phosphorylated Nedd4-2 were detected using rabbit antibodies anti-NEDD4L (4013S), anti-p-NEDD4L-S448 (8063S) and anti-p-NEDD4L-S342 (12146S) (Cell Signaling). Total and phosphorylated glycogen synthase kinase 3 (GSK3-β) were detected using rabbit antibodies anti-GSK3-*β* (9315) and anti-pGSK3α/*β*-Ser21/9 (9331) (Cell Signaling). Chemiluminescence signals were analyzed using Image Lab software 6.0 (Bio-Rad).

### EEG recordings and analysis

Cortical and hippocampal electroencephalogram (EEG) activity was recorded as previously described (40). B6.Tg.sgk1 or B6.WT anesthesized mice (1.5% isoflurane in 100% oxygen) were implanted a nickel-chromium electrode (140 μm) in the prefrontal cortex (1.5 mm rostral, 1,5 mm lateral and 1 mm ventral to bregma), a second electrode in the CA1 region of hippocampus (2.4 mm rostral, 1.5 mm lateral, and 1.5 mm ventral to bregma) and two stainless steel for ground and indifferent references. Recovering took place in the home cages in a sound attenuated chamber under 12:12 h light-dark regime, constant temperature (22°C), and *ad libitum* access to food and water. Nine days after surgery, mice were transferred to a circular cage and the implanted cap fixed to a rotating anti-gravitational connector allowing free movement. After a period of habituation of 72 h, 24 h of uninterrupted recording were acquired starting at least 1 h before KA injection. EEG signals of the cortex and hippocampus were filtered (0.5-200 Hz), amplified (x5000-10000), and digitized at 1 kHz (CyberAmp 380, Digidata 1440A, AxoScope 10 software, Molecular Devices). EEG synchronized video recordings (Logitech Software and Cerberus 4.0 Beta) were used to score behavioral Racine stages.

Raw EEG recordings were analyzed using Spike2 V5 software and custom scripts in MATLAB v.5.3 (MathWorks). First, all data were normalized using the z-score to allow for EEG comparisons. EEG segments that contained epileptic activity were determined by time-frequency analysis, which was performed using the Fast Fourier Transform (FFT) to calculate the power spectrum density (0- 150 Hz) for 5 s windows (50% overlapping, Hamming function) with 0.20 Hz resolution. Root-Mean Square (RMS) power of the signals was calculated and smoothed (5-s window) to obtain mean and standard deviation of the normalized amplitude of the cortex and hippocampal signal. Generalized seizures (GS) were defined as episodes of spike discharges of high amplitude (≥2x average baseline amplitude) and high frequency (>30 Hz), and detected automatically. Further analysis rendered values for latency, number and duration of GSs, and inter-seizure interval.

### Gene expression analysis

Quantitative real-time PCR (qPCR) was performed to evaluate relative expression of *Sgk1* isoforms (SGK1 and SGK1.1), KCNQ2, KCNQ3, and KA receptors GRIK 1-5 in B6.Tg.sgk1 and B6.WT mice. RNA extraction was performed from snap-frozen hippocampal and neocortex tissues from mice, using a commercial kit (Real Total RNA spin plus). Quantity and purity of RNA was measured with a Nanodrop 1000 (Thermo Scientific). Equal amounts of total RNA were retrotranscribed to cDNA with iScript cDNA synthesis kit (BioRad). Each qPCR reaction was performed in triplicate using specific primers (**Supplementary Table 1**) and iQ SYBR Green Supermix (Bio-Rad) in a CFX96 Touch™ Real-Time PCR Detection System (Bio-Rad). Relative quantification of mRNA abundance was performed using Biogazelle qBASE plus-2 software. Data are average fold-change ± SEM.

### Statistics

Statistical analysis was performed using Prism7 (GraphPad) and SigmaPlot (SigmaPlot) as specified in the figure legends. Statistical significance was set at p < 0.05. Data are mean ± SEM.

## Acknowledgments

We thank Dr. Teresa Acosta Almeida for help with gene expression data analysis and Dr. Ricardo Gómez for useful comments on the manuscript. This work was funded by grants BFU2015-66490- R, BFU2015-71078-P and BFU2014-58999-P (Ministerio de Economia y Competitividad [MINECO], Spain) to T.G., L.C.B. and J.A.L, respectively. D.A.R., L.C.B., J.A.L. and T.G. are members of the Red de Excelencia “Iniciativa Española en Canales Iónicos” (BFU2015-70067- REDC, MINECO, Spain). T.G. was supported by Programa Ramón y Cajal (RYC-2012-11349, MINECO, Spain). N.A.C. was supported by a predoctoral fellowship from Gobierno de Canarias (Agencia Canaria de Investigación, Innovación y Sociedad de la Información, ACIISI, TESIS20120160). E.M.B. is supported by a F.P.I. predoctoral Fellowship (BES-2016-077337) from MINECO, Spain. The authors declare no competing interests.

## SUPPLEMENTAL DATA

**Supplemental Figure 1.**
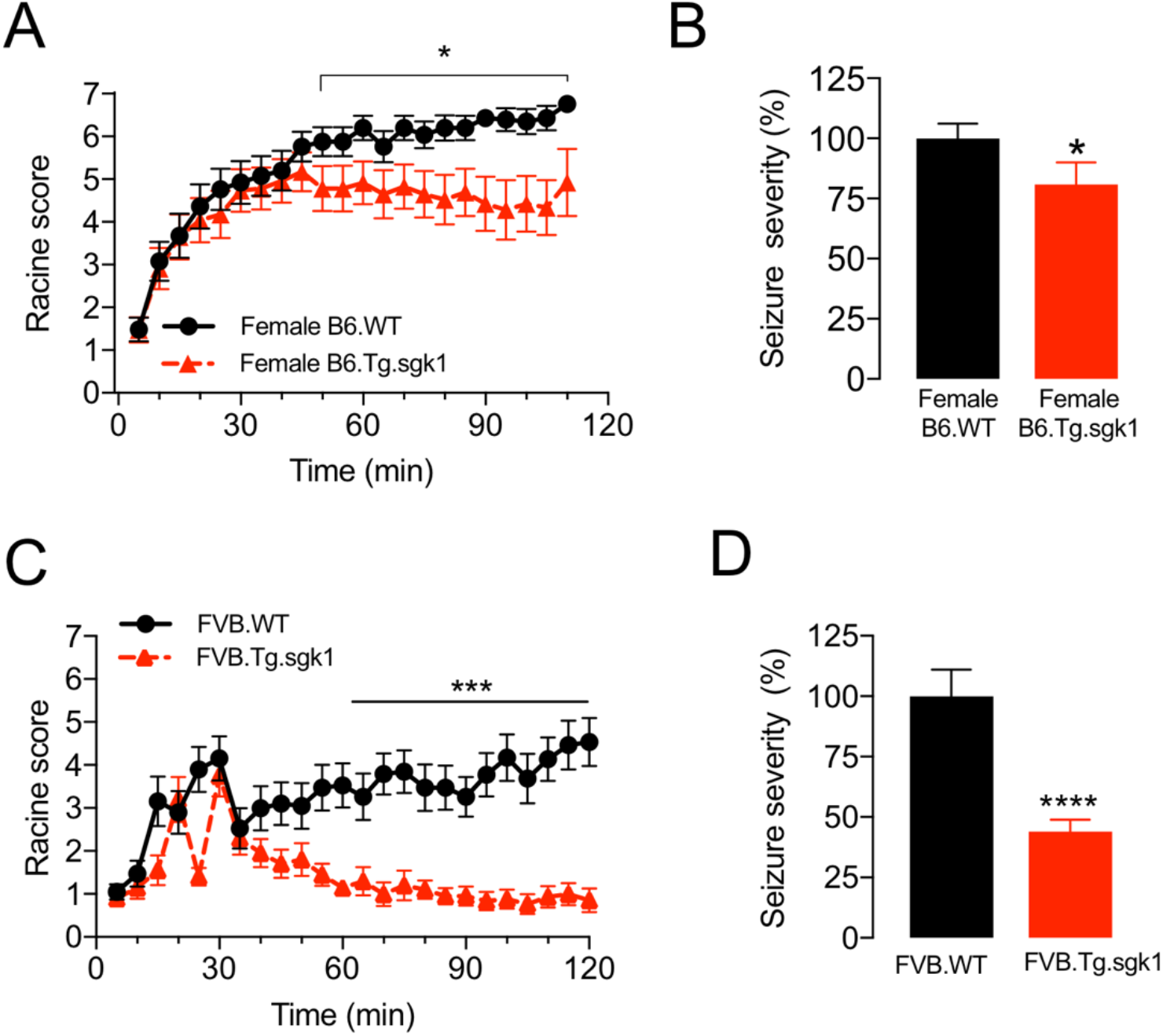
Seizure protection associated to *Sgk1* activation is independent of sex and genetic background. (**A)** Averaged Racine score measured at 5 min-intervals for female mice of both genotypes. Data are mean ± SEM; * p < 0.05, ** p < 0.005, *two-tailed Student t-test*. The evolution of seizure-like behavior in female mice, including the differences between genotypes, was similar to that observed in males. Of note, our results showed a significantly higher percentage of female animals experiencing severe seizures as compared to their male counterparts for both B6.WT and B6.Tg.sgk1 genotypes (61% B6.Tg.sgk1 females reaching tonic-clonic seizures *vs.* 92% B6.WT), in agreement with previous findings indicating higher susceptibility of females to seizures ((20) but see also (21)). (**B)** Integrated seizure severity corresponding to female mice, represented as mean ± SEM; * p < 0.5, *Two-tail, Student t-test*. (**C)** Time-course analysis of the Racine score for 5 min intervals in Tg.sgk1 mice backcrossed into a FVB/NJ background, which has been proposed to be more susceptible to KA-induced seizures (41); *** p < 0.0005, *two-tailed Student t-test*. (**D)** Integrated seizure severity of FVB.Tg.sgk1 mice compared to FVB.WT. Note that transgenic mice showed a regression towards less severe stages after the initial period of seizure progression, while FVB.WT mice continued progressing towards tonic-clonic seizure levels over at least 1 h. Data are mean ± SEM; **** p = 0.0001, *two-tailed Student t-test*.

**Supplemental Figure 2.**
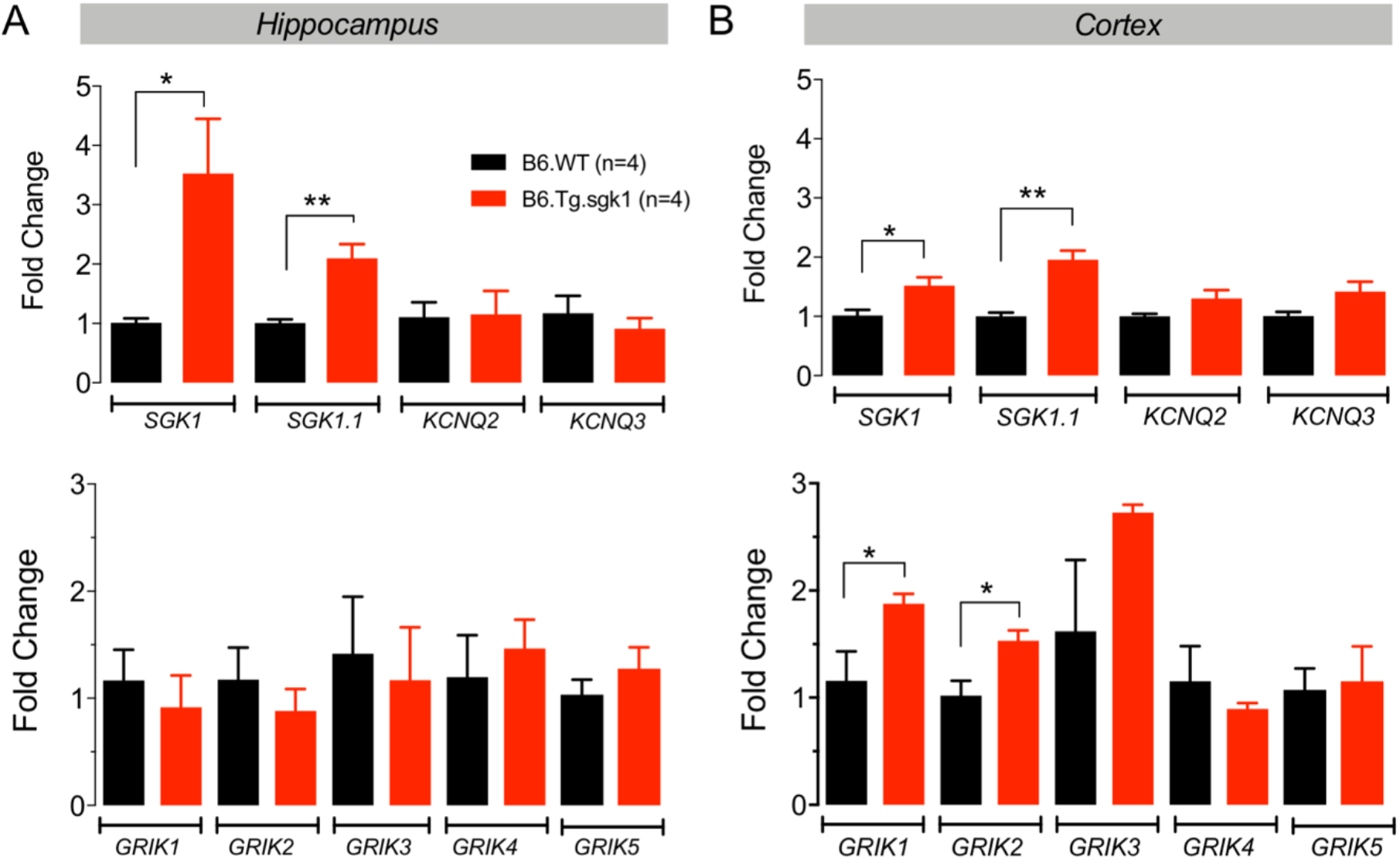
Analysis of mRNA expression of SGK1, SGK1.1, KCNQ2 and KCNQ3 (*top graphs*), KA receptors GRIK1 (GluR5), GRIK2 (GluR6), GRIK3 (GluR7), GRIK4 (KA1) and GRIK5 (KA2) (*Bottom graphs*) in **(A)** hippocampus and **(B)** cortex. Note that basal expression levels of SGK1 and SGK1.1 were significantly increased in B6.Tg.sgk1 hippocampus and cortex. No variations in KCNQ2/3 gene expression were observed between genotypes. No significant alterations in GRIK1-5 KA receptor expression were observed in hippocampus, although slightly increased expression of GRIK1 and GRIK2 was noticed in the cortex of transgenic animals. Data were normalized using the relative expression of two housekeeper genes (RPL13a and CycA). Bars represent fold change ± SEM in reference to B6.WT. * p < 0.05, ** p < 0.005, *two-tailed Student t-test*.

**Supplemental Figure 3.**
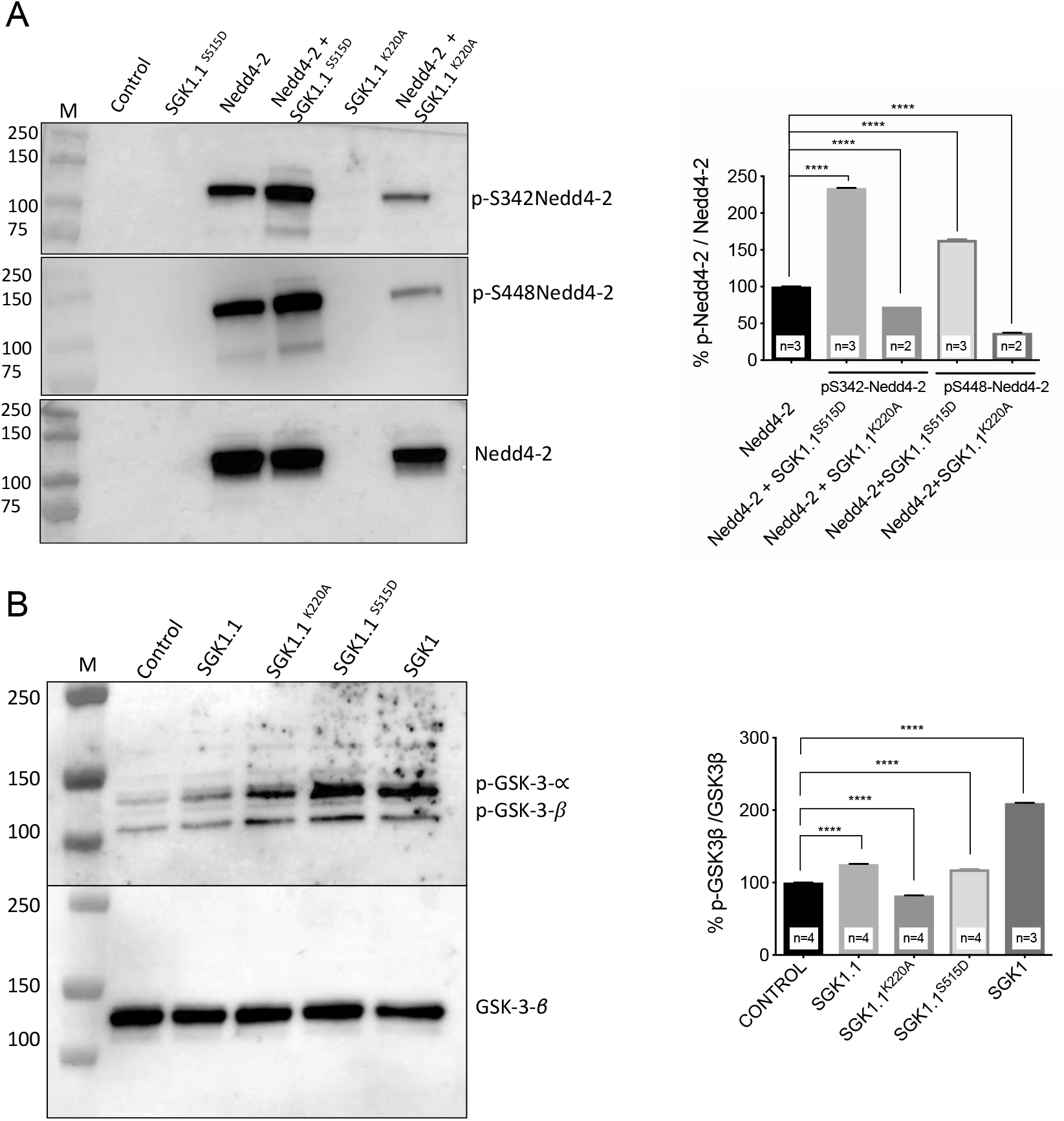
SGK1.1 phosphorylates Nedd4-2 and GSK3−β in heterologous expression systems. **(A)** *Left*, representative immunoblot of protein extracts from HEK293 cells transiently co-transfected with the indicated combinations of Nedd4-2, SGK1.1^S515D^ (constitutively active SGK1.1 mutant) and SGK1.1^K220A^ (kinase inactive mutant). Top panel, labeling with pS342- Nedd4-2 antibody. Middle panel, pS448-Nedd4-2. Total Nedd4-2 was used as internal standard. Untransfected cells were used as negative control. M, molecular mass markers (in kDa). *Right*: Quantification of pS342-Nedd4-2 and pS448-Nedd4-2 levels relative to levels of total Nedd4-2. Bars are mean ± SEM; ****p < 0,0001, *one-way ANOVA, Dunnett’s multiple comparisons test*. **(B)** *Left:* Representative immunoblot of HeLa cells transfected with the indicated plasmid combinations and probed with an antibody against pS21-GSK3-*α* and pS9-GSK3*−β*. Total GSK3*−β* was used as internal standard. Untransfected cells were used as negative control. *Right*: Quantification of pS9- GSK3-*β* levels relative to total GSK3-*β*. Bars represent mean ± SEM; ****p < 0,0001, *one-way ANOVA, Dunnett’s multiple comparisons test*.

**Supplemental Figure 4.**
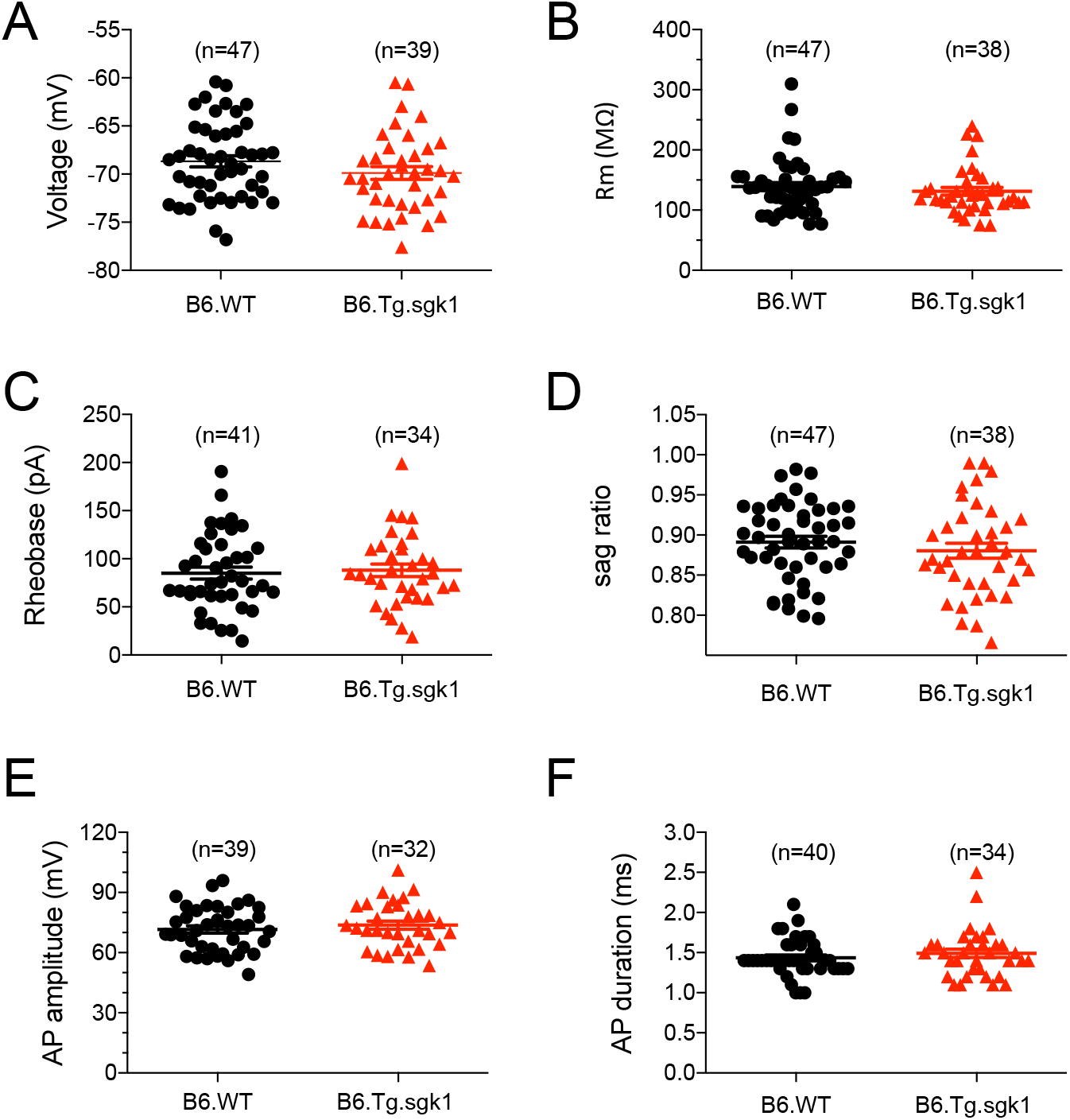
Membrane properties and action potential characteristics of B6.WT and B6.Tg.sgk1 CA1-PN. **(A)** resting membrane potential, **(B)** membrane resistance, **(C)** rheobase, sag ratio, **(E)** action potential amplitude, and **(F)** action potential duration. No significant statistical differences were observed between genotypes for any of these parameters (*two-tailed Student t-tests*).

**Supplementary Table 1.**
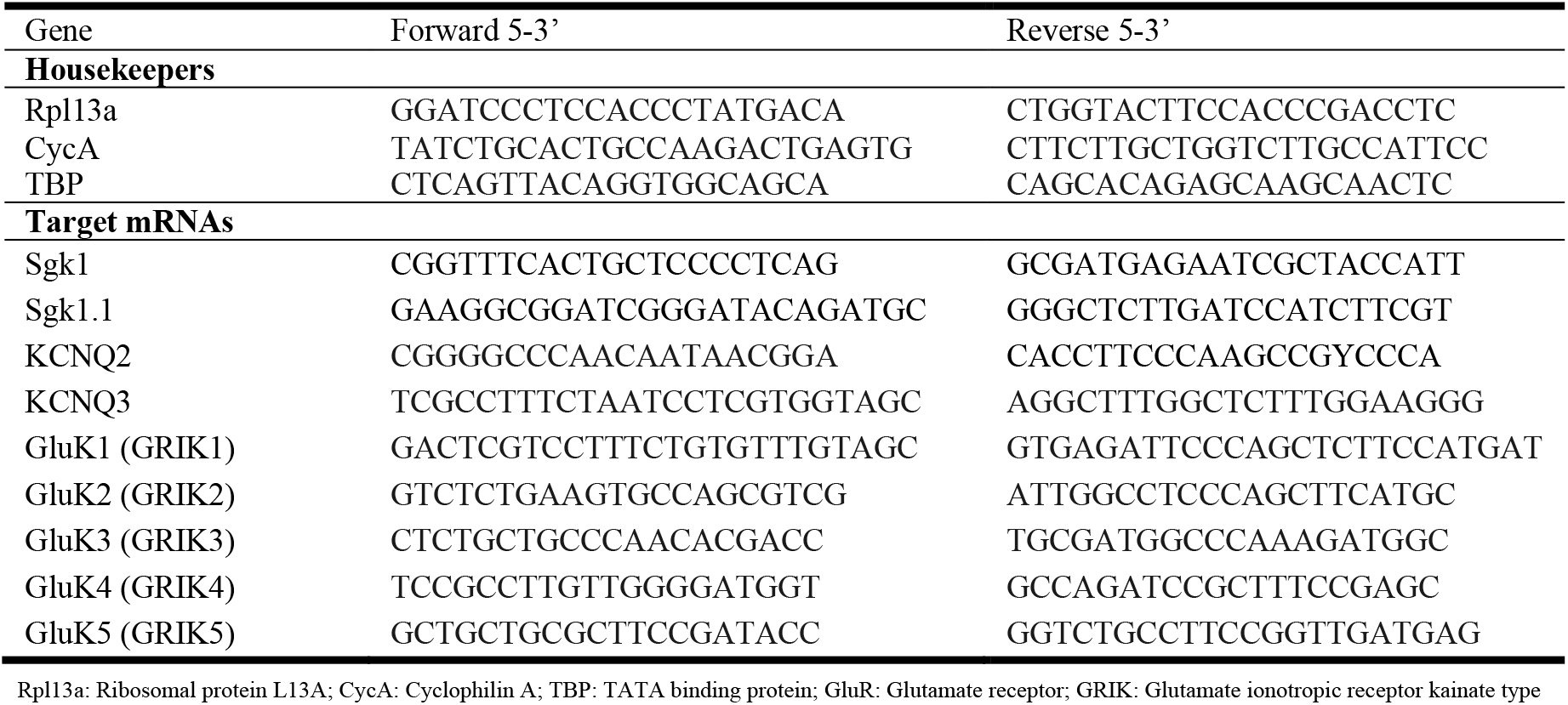
Primers pairs used for qPCR analysis of mRNA expression.

